# Uncovering the heterogeneity and temporal complexity of neurodegenerative diseases with Subtype and Stage Inference

**DOI:** 10.1101/236604

**Authors:** Alexandra L Young, Razvan V Marinescu, Neil P Oxtoby, Martina Bocchetta, Keir Yong, Nicholas Firth, David M Cash, David L Thomas, Katrina M Dick, Jorge Cardoso, John van Swieten, Barbara Borroni, Daniela Galimberti, Mario Masellis, Maria Carmela Tartaglia, James B Rowe, Caroline Graff, Fabrizio Tagliavini, Giovanni Frisoni, Robert Laforce, Elizabeth Finger, Alexandre Mendonça, Sandro Sorbi, Jason D Warren, Sebastian Crutch, Nick C Fox, Sebastien Ourselin, Jonathan M Schott, Jonathan D Rohrer, Daniel C Alexander on behalf of the Genetic FTD Initiative, GENFI and the Alzheimer’s Disease Neuroimaging Initiative

## Abstract

The heterogeneity of neurodegenerative diseases is a key confound to disease understanding and treatment development, as study cohorts typically include multiple phenotypes on distinct disease trajectories. Here we present a new machine learning technique – Subtype and Stage Inference (SuStaIn) – able to uncover data-driven disease phenotypes with distinct temporal progression patterns, from widely available crosssectional patient studies. Results from imaging studies in two neurodegenerative diseases reveal new subgroups and their distinct trajectories of regional neurodegeneration. In genetic frontotemporal dementia, SuStaIn identifies genotypes from imaging alone, validating its ability to identify subtypes, and characterises within-group heterogeneity for the first time. In Alzheimer’s disease, SuStaIn uncovers three subtypes, uniquely revealing their temporal complexity. SuStaIn provides fine-grained patient stratification, which substantially enhances the ability to predict conversion between diagnostic categories over standard models that ignore subtype (p=7.18×10-^-4^) or temporal stage (p=3.96×10^−5^). SuStaIn thus offers new promise for enabling disease subtype discovery and precision medicine.

## Introduction

Neurodegenerative disorders, such as frontotemporal dementia (FTD) and Alzheimer’s disease (AD), are biologically heterogeneous, producing high variance in *in vivo* disease biomarkers, such as volumetric measurements from imaging, protein measurements from lumbar puncture, or behavioural measurements from psychometrics, which reduces their utility in disease studies and management. Key contributors to this heterogeneity are that individuals belong to a range of disease subtypes (giving rise to *phenotypic heterogeneity*) and are at different stages of a dynamic disease process (producing *temporal heterogeneity*). Previous studies aiming to explain biomarker variance typically focus on a single aspect of this heterogeneity: phenotypic heterogeneity at a coarse, typically late, disease stage, or temporal heterogeneity in a broad population. However, the inability to disentangle the range of subtypes from the development and progression of each over time limits the biological insight these techniques can provide, as well as their utility for patient stratification. Constructing a comprehensive picture separating phenotypic and temporal heterogeneity, i.e. identifying distinct subtypes and characterising the development and progression of each remains a major current challenge. However, such a picture would provide novel insights into underlying disease mechanisms, and enable accurate fine-grained patient stratification and prognostication, facilitating precision medicine in clinical trials and healthcare.

Both FTD and AD exhibit substantial pathologic, genetic, and clinical heterogeneity. In FTD a large proportion of cases (around a third) are inherited on an autosomal dominant basis, with mutations in progranulin (*GRN*), microtubule-associated protein tau (*MAPT*) and chromosome 9 open reading frame 72 (*C9orf72*) being the most common causes. Of the major genetic groups, *GRN* mutations are associated with TDP-43 type A pathology, *MAPT* mutations with tau inclusions, and expansions in *C9orf72* with type A or type B TDP-43 pathology^1^. AD instead has a single pathological characterisation: the presence of both amyloid plaques and neurofibrillary tangles, and the proportion of autosomal dominant cases is much smaller, accounting for between 1% and 6% of cases^2^. The pathological heterogeneity observed in AD consists of variation in the distribution of neurofibrillary tangles, with 25% of patients having an atypical distribution of neurofibrillary tangles (described as hippocampal-sparing or limbic-predominant) on autopsy at the time of death^3^. Both FTD and AD exhibit a diverse range of clinical syndromes. FTD has both behavioural and language presentations, and in genetic FTD the clinical syndromes can further include atypical parkinsonism and amytrophic lateral sclerosis (ALS). In AD, the major clinical syndrome is broadly divided into amnestic and rarer non-amnestic variants, with non-amnestic variants including language variant AD, lopogenic progressive aphasia, visuoperceptive variant AD, posterior cortical atrophy (PCA), and frontal variant AD^4^.

Previous studies of neurodegenerative disease heterogeneity have focussed on either temporal heterogeneity (i.e. subjects appear different at different disease stages) or phenotypic heterogeneity (i.e. distinct groups of subjects appear different even at the same disease stage), but rarely both. We refer to these two approaches as *stages-only models*, which account for temporal heterogeneity but not phenotypic heterogeneity, and *subtypes-only models*, which account for phenotypic heterogeneity but not temporal heterogeneity. Stages-only models arise for example from regression against disease stage^5,6^, and data-driven disease progression modelling^7-15^. Although such models have provided new understanding of the temporal progression of a range of conditions, the inherent assumption that all individuals have a single phenotype, i.e. follow approximately the same trajectory, is a key limitation. At best, this limits the biological insight and the accuracy of stratification they can provide, but potentially could also lead to erroneous conclusions. Subtypes-only models use, for example, clustering (e.g. ^16-23^) to identify distinct groups, or group individuals using information independent of the model, such as genetics (e.g. ^24^) or post-mortem examination (e.g. ^25-28^) for models based on in-vivo imaging. With typical subtypes-only models, the limitation is the inherent assumption that all subjects are at a common disease stage so that the cohort has no temporal heterogeneity. This requires *a priori* staging and selection of individuals, which is typically crude in practice leaving models that are not specific to subtype differences. Models of both disease subtype and stage heterogeneity have been constructed previously for the small proportion of neurodegenerative diseases that are inherited on an autosomal-dominant basis. For example, Rohrer et al^29^ investigate temporal heterogeneity within genetic groups by regressing imaging markers against an estimated age of onset (from family history). However, such studies lack the ability to identify novel within-genotype phenotypes, and the temporal resolution of the recovered genotype progression patterns is limited by inaccuracy of the a-priori staging.

This paper presents Subtype and Stage Inference (SuStaIn) (see conceptual overview in Figure 1): a computational technique that disentangles temporal and phenotypic heterogeneity to identify population subgroups with common patterns of disease progression. SuStaIn is a new unsupervised machine-learning technique that uniquely builds on and combines ideas from clustering (e.g. ^16-23^) and data-driven disease progression modelling (e.g. ^7-10^,^12^). The combination uniquely enables SuStaIn to group individuals with common phenotypes across the range of disease stages. It determines the number of subtypes that the available data can support, reconstructs the trajectory of stages within each subtype, and assigns a probability of each subtype and stage to each subject. These features offer the potential for important new insights into disease biology, e.g. by revealing the earliest sites of disease and subsequent spreading patterns and thus supporting mechanistic models (e.g. ^30,31^) of disease aetiology without the confounds of phenotypic heterogeneity. They also provide a mechanism for *in vivo* fine-grained stratification at early disease stages, facilitating precision medicine.

**Figure 1.**
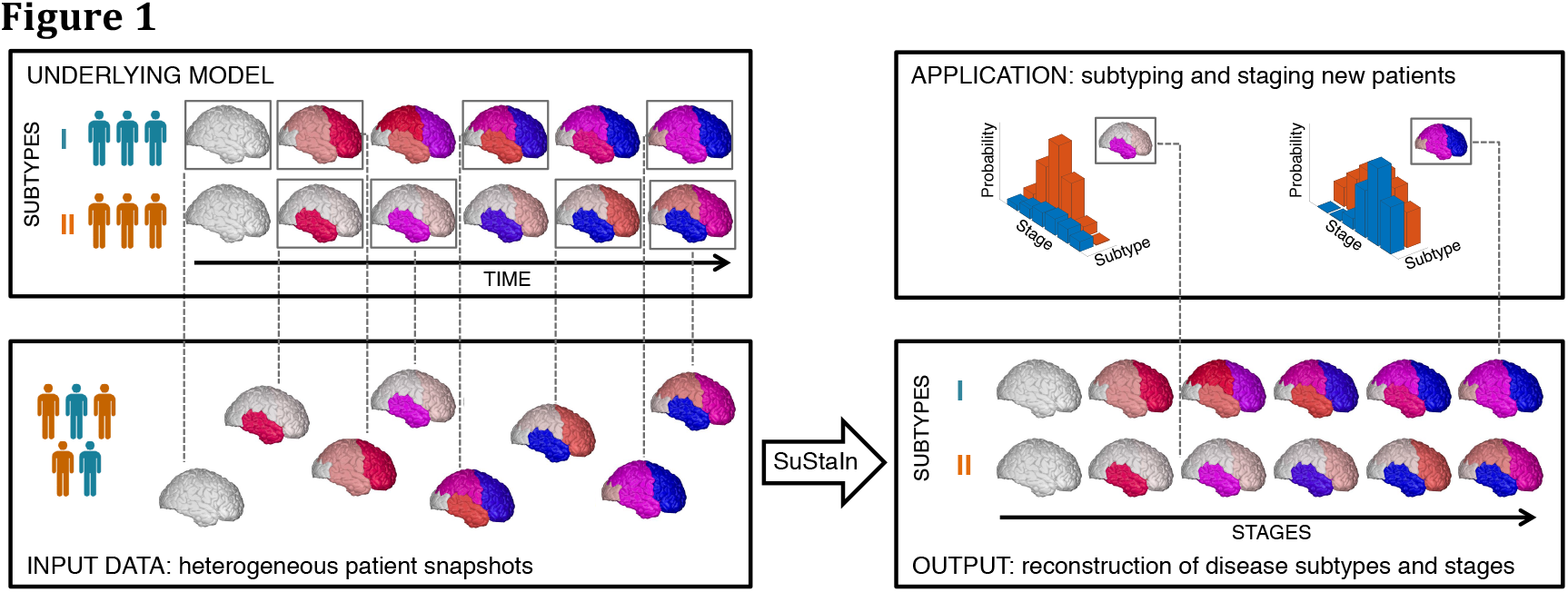
Conceptual overview of SuStaIn. The Underlying Model panel (top left) considers a patient cohort to consist of an unknown set of disease subtypes. The input data (Input Data panel, bottom left), which can be entirely cross-sectional, contains snapshots of biomarker measurements from each subject with unknown subtype and unknown temporal stage. SuStaIn recovers the set of disease subtypes and their temporal progression (as shown in Output panel, bottom right) via simultaneous clustering and disease progression modelling. Given a new snapshot, SuStaIn can estimate the probability the subject belongs to each subtype and stage, by comparing the snapshot with the reconstruction (as shown in Application panel, top right). In this figure two hypothetical disease subtypes are modelled, labelled I and II, and the biomarkers are regional brain volumes, but SuStaIn is readily applicable to any scalar disease biomarker. The colour of each region indicates the amount of pathology in that region, ranging from white (no pathology) to red to magenta to blue (maximum pathology).

Here, we use SuStaIn with structural magnetic resonance imaging (MRI) data sets from cohorts of genetic FTD and AD patients. In each case, SuStaIn provides a novel data-driven taxonomy (set of subtypes and stages), as well as new and detailed pictures of the progression of neurodegeneration within each of the data-driven subgroups. The genetic FTD data set provides a validation of SuStaIn’s ability to identify subgroups with distinct temporal progression patterns, as the different genotypes are known to have distinct patterns of neurodegeneration visible as brain atrophy in MRI ^29^. SuStaIn identifies subtypes from imaging alone that map closely onto the genotypes and reconstructs patterns of neurodegeneration that reflect analysis of the individual genetic groups. It further uncovers two distinct previously unseen within-genotype phenotypes for carriers of a mutation in the *C9orf72* gene, while finding the *MAPT* and *GRN* mutation groups are more homogeneous. In AD, SuStaIn identifies three distinct subtypes and reconstructs their previously unseen temporal progression. In both neurodegenerative diseases, we demonstrate strong identifiability of the SuStaIn subtypes, i.e. we can assign patients to subtype, which subtypes-only models in the literature are unable to do; see e.g. ^23^. Even at very early disease stages, at least a proportion of individuals show strong alignment with particular subtypes, which highlights the potential utility in precision medicine. In AD, we show that SuStaIn subtype and stage enhance the ability to predict conversion between diagnostic categories substantially beyond subtypes-only or stages-only models.

## Results

### Subtype progression patterns

We demonstrate SuStaIn in two neurodegenerative diseases, genetic FTD and sporadic AD, using cross-sectional regional brain volumes from MRI data in the GENetic Frontotemporal dementia Initiative (GENFI) and the Alzheimer’s Disease Neuroimaging Initiative (ADNI). GENFI investigates biomarker changes in carriers of mutations in progranulin (GRN), microtubule-associated protein tau (*MAPT*), and chromosome 9 open reading frame 72 (*C9orf72*) genes, which cause FTD. *GRN* and *MAPT* mutations are known to be associated with more distinct phenotypes, whereas *C9orf72* is a heterogeneous group. We used GENFI as a test data set with a partially known ground truth partly for validation, as we expect SuStaIn to identify genetic groups as distinct phenotypic subtypes, but also to investigate the phenotypic and temporal heterogeneity within genotypes. Specifically, we tested SuStaIn on the combined data set from all mutation carriers in GENFI (Figure 2A), without using knowledge of their genotype, and compared the resulting subtype progression patterns with (a) participant’s genotype labels (Figure 2B), and (b) subtype progression patterns obtained from each genetic type separately (Figure S1). Next, we used SuStaIn to identify sporadic AD subtypes from ADNI and characterise their progression from early to late disease stages (Figure 3). We tested consistency of the SuStaIn subtypes in a largely independent dataset – ADNI 1.5T MRI scans rather than the main 3T data set (Figure 4). In each disease we cross-validated our results to test the reproducibility of the subtypes and estimated progression patterns (Figure S2).

**Figure 2.**
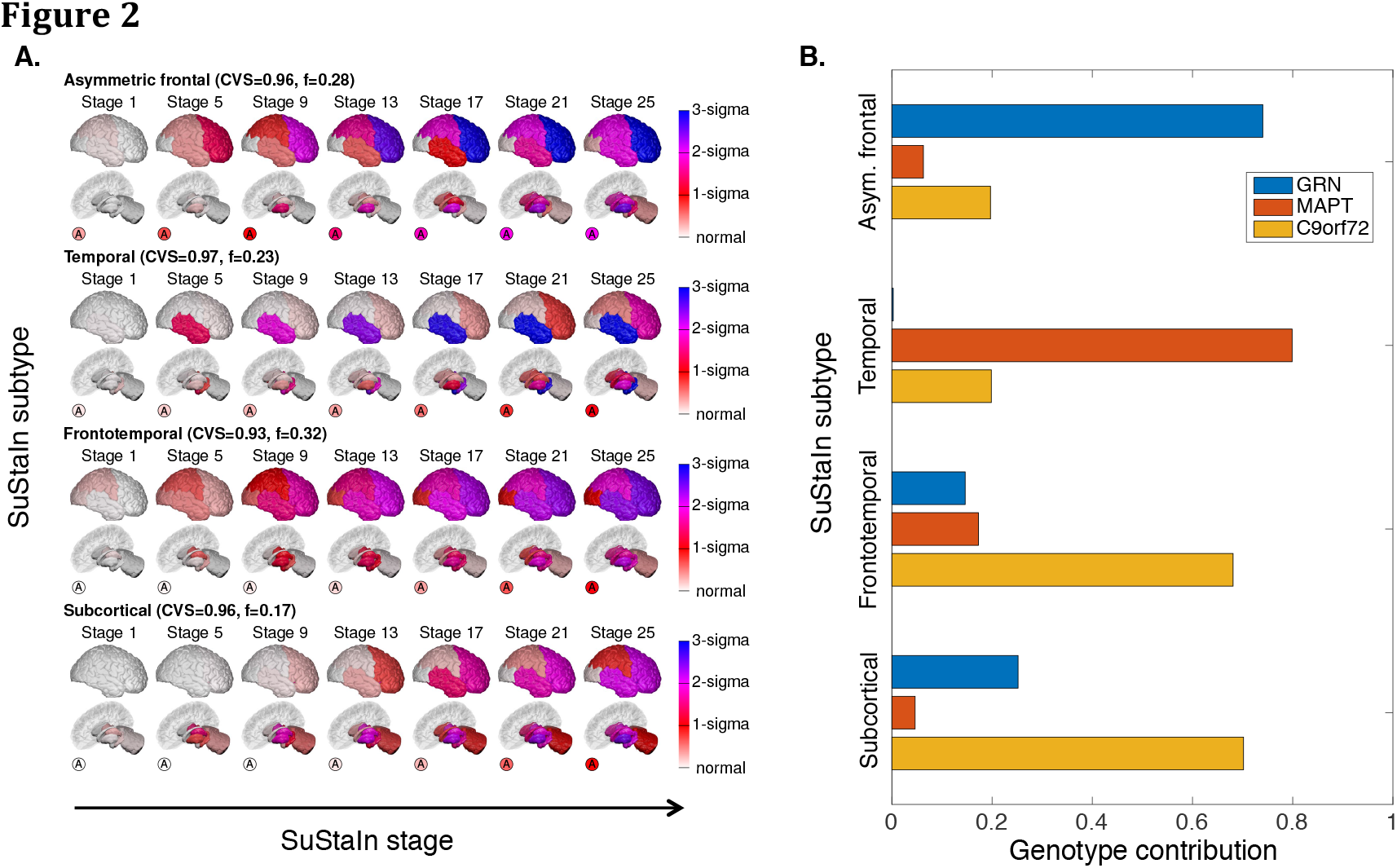
SuStaIn modelling of genetic frontotemporal dementia using GENFI data. Subfigure A shows the progression pattern of each of four subtypes that SuStaIn identifies. Each progression pattern consists of a sequence of stages in which regional brain volumes in mutation carriers (symptomatic and presymptomatic) reach different z-scores relative to non-carriers. Intuitively (for a more precise description see Methods: Uncertainty Estimation), at each stage the colour in each region indicates the level of severity of volume loss: white is unaffected; red is mildly affected (z-score of 1); magenta is moderately affected (z-score of 2); and blue is severely affected (z-score of 3 or more). The circle labelled ‘A’ indicates the asymmetry of the atrophy pattern (absolute value of the difference in volume between the left and right hemispheres divided by the total volume of the left and right hemispheres) at each stage for each subtype. CVS is the model cross-validation similarity (see Methods: Similarity between two progression patterns): the average similarity of the subtype progression patterns across cross-validation folds, which ranges from 0 (no similarity) to 1 (maximum similarity), f is the proportion of participants estimated to belong to each subtype. Subfigure B shows the contribution of each genotype to each of the SuStaIn subtypes. This is calculated as the probability an individual has a particular genotype given that they belong to a particular subtype.

**Figure 3.**
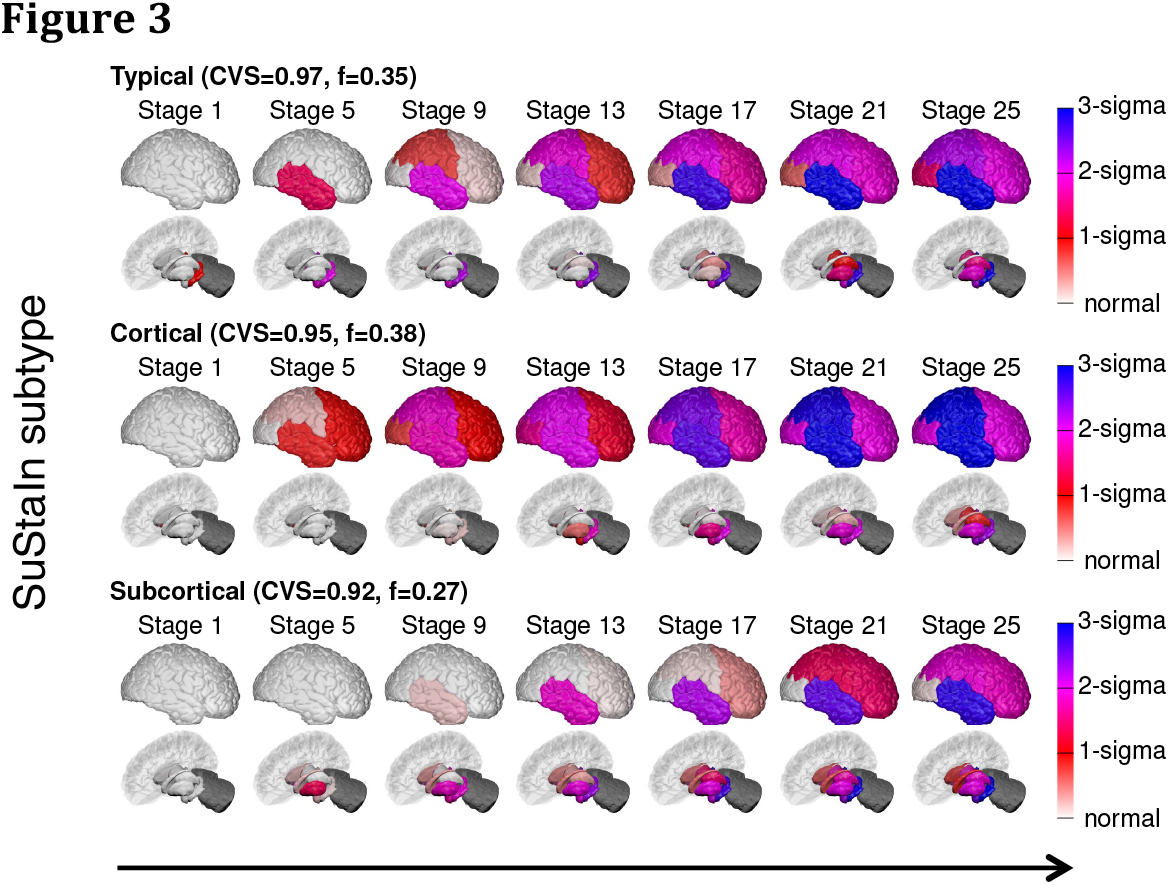
SuStaIn modelling of sporadic Alzheimer’s disease using ADNI data. Each row shows the progression pattern of each of the three subtypes identified by SuStaIn. Diagrams as in Figure 2, but the z-scores are measured relative to amyloid-negative (cerebrospinal fluid (CSF) Aβ1-42>192pg/ml) cognitively normal subjects, i.e. cognitively normal subjects with no evidence of amyloid pathology on CSF. The cerebellum was not included as a region in the Alzheimer’s disease analysis and so is shaded in dark grey.

**Figure 4.**
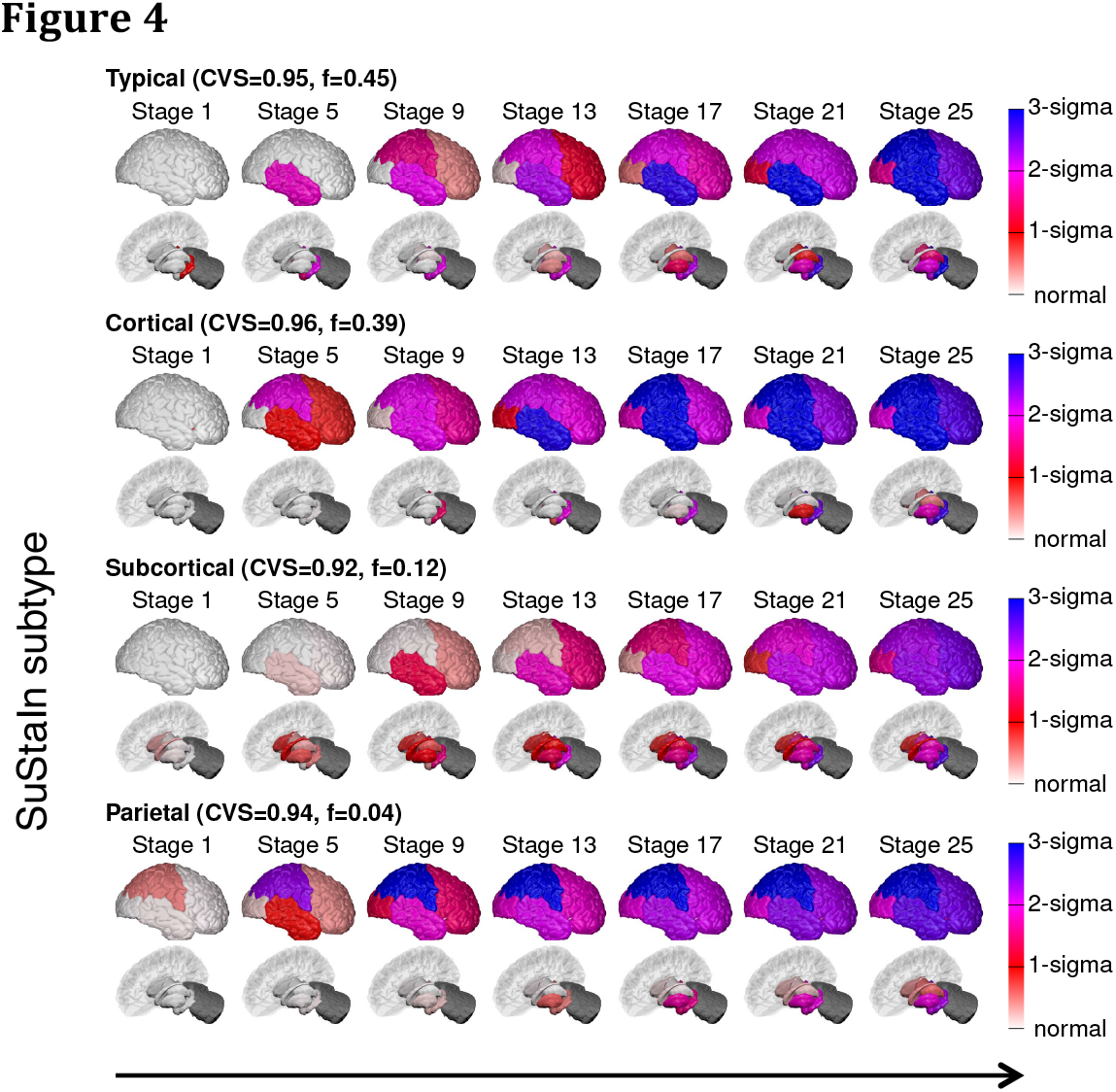
Reproducibility of the SuStaIn subtypes in Figure 3 in a largely independent Alzheimer’s disease dataset (only 59 subjects are in both the 576 subject dataset used to generate this figure and the 793 subject dataset used in Figure 3) consisting of those with regional brain volume measurements from 1.5T MRI scans, rather than 3T MRI scans. Diagrams are as in Figure 3, with each row showing the progression pattern of each of the subtypes identified by SuStaIn. SuStaIn modelling identifies three major subtypes: a typical, a cortical and a subcortical subtype, which are in good agreement with the three subtypes in Figure 3, as well as an additional very small outlier group (only 4%) with a subtype we term “parietal”. This small subgroup may represent outliers with a posterior cortical atrophy phenotype.

#### SuStaIn reveals novel within genotype phenotypes in FTD

Figure 2 shows that SuStaIn successfully identifies the progression patterns of the different genetic groups in GENFI, without prior knowledge of genotype, and further reveals that the *C9orf72* group is phenotypically heterogeneous, finding two neuroanatomical subtypes. Figure 2A shows the four subtypes that SuStaIn finds from the full set of all mutation carriers in GENFI. We refer to them as the asymmetric frontal lobe subtype, temporal lobe subtype, frontotemporal lobe subtype, and subcortical subtype. Figure 2B reveals that *GRN* mutation carriers are the main contributors to the asymmetric frontal lobe subtype, *MAPT* mutation carriers are the main contributors to the temporal lobe subtype, and *C9orf72* mutation carriers are the main contributors to both the frontotemporal lobe subtype and the subcortical subtype. This suggests that there are two distinct subtypes in the *C9orf72* group. Application of SuStaIn to each genetic type separately supports this finding by demonstrating that the *GRN* mutation carriers are best described as a single asymmetric frontal lobe subtype, the *MAPT* mutation carriers are best described as a temporal lobe subtype and the *C9orf72* mutation carriers are best described as two distinct disease subtypes: a frontotemporal lobe subtype and a subcortical subtype. SuStaIn additionally finds an outlier cluster in the *MAPT* group for which the progression pattern has high uncertainty. This high uncertainty likely prevents the cluster from being detected when applying SuStaIn to all mutation carriers in Figure 2 as this small proportion of outliers can be sufficiently modelled by the three alternative subtype progression patterns. Figure S1 shows that the subtype progression patterns for each genetic type are in good agreement with those found in the full set of all mutation carriers (Figure 2A). Figure S2A shows that the four subtypes estimated in Figure 2A are reproducible under cross-validation, with a high average similarity between cross-validation folds of greater than 93% for each subtype. Altogether these results provide strong evidence that the *C9orf72* group are phenotypically heterogeneous, expressing two distinct subtypes, whereas the *GRN* and *MAPT* groups express more homogeneous phenotypes.

#### SuStaIn identifies three subtype progression patterns in AD

Figure 3 shows the temporal progression of the three neuroanatomical subtypes that SuStaIn identifies from ADNI, which we term “typical”, “cortical” and “subcortical”. SuStaIn reveals that for the typical subtype atrophy starts in the hippocampus and amygdala; for the cortical subtype in the nucleus accumbens, insula and cingulate; and for the subcortical subtype in the pallidum, putamen, nucleus accumbens and caudate. Figure S2B shows that these three subtypes are reproducible under cross-validation, giving an average similarity between cross-validation folds of greater than 92% for each subtype.

#### AD subtype progression patterns are reproducible in an independent dataset

Figure 4 shows that the three subtypes in Figure 3 are reproducible in a largely independent dataset consisting of regional brain volumes derived from 1.5T rather than 3T MRI scans. From the 1.5T data, SuStaIn broadly replicates the three major clusters found in the 3T data, again finding a typical, cortical and subcortical subtype. The origin of atrophy for each subtype is in general agreement with the 3T data: atrophy begins in the hippocampus and amygdala for the typical subtype, in the insula and cingulate for the cortical subtype; and in the pallidum, putamen and caudate for the subcortical subtype. The main difference compared to the 3T data is that the nucleus accumbens is not indicated as an early region to atrophy in the 1.5T data for the cortical and subcortical subtypes. SuStaIn additionally identifies a small proportion (4%) of outliers with a parietal subtype in the 1.5T data.

### Disease subtyping and staging

We investigated SuStaIn’s capability for reliable stratification in each neurodegenerative disease (Figure 5) to determine the potential for homogeneous cohort identification. First, we assessed how reliably SuStaIn assigns patients to subtypes (Figure 5A and B). Specifically, in genetic FTD, we tested the consistency of SuStaIn subtypes with the different genetic types in symptomatic mutation carriers. We also compared the discriminative power of the SuStaIn subtypes against a subtypes-only model, i.e. clustering without progression modelling: a comparable model that accounts for phenotypic heterogeneity but not temporal heterogeneity (Figure 6 and Table 1). Second, we assessed the reliability of the SuStaIn stages in each disease (Figure 5C and D) by comparison with clinical diagnostic categories. In ADNI, where clinical follow-up information is available, we further examined the ability of SuStaIn subtypes and stages to predict relevant outcomes, by determining whether SuStaIn subtype and/or stage modify the risk of conversion between diagnostic categories (Table 2). We compared the predictive power with a subtypes-only model, and stages-only model, i.e. progression modelling without clustering: a comparable model that accounts for temporal heterogeneity but not phenotypic heterogeneity (Table S1).

**Figure 5.**
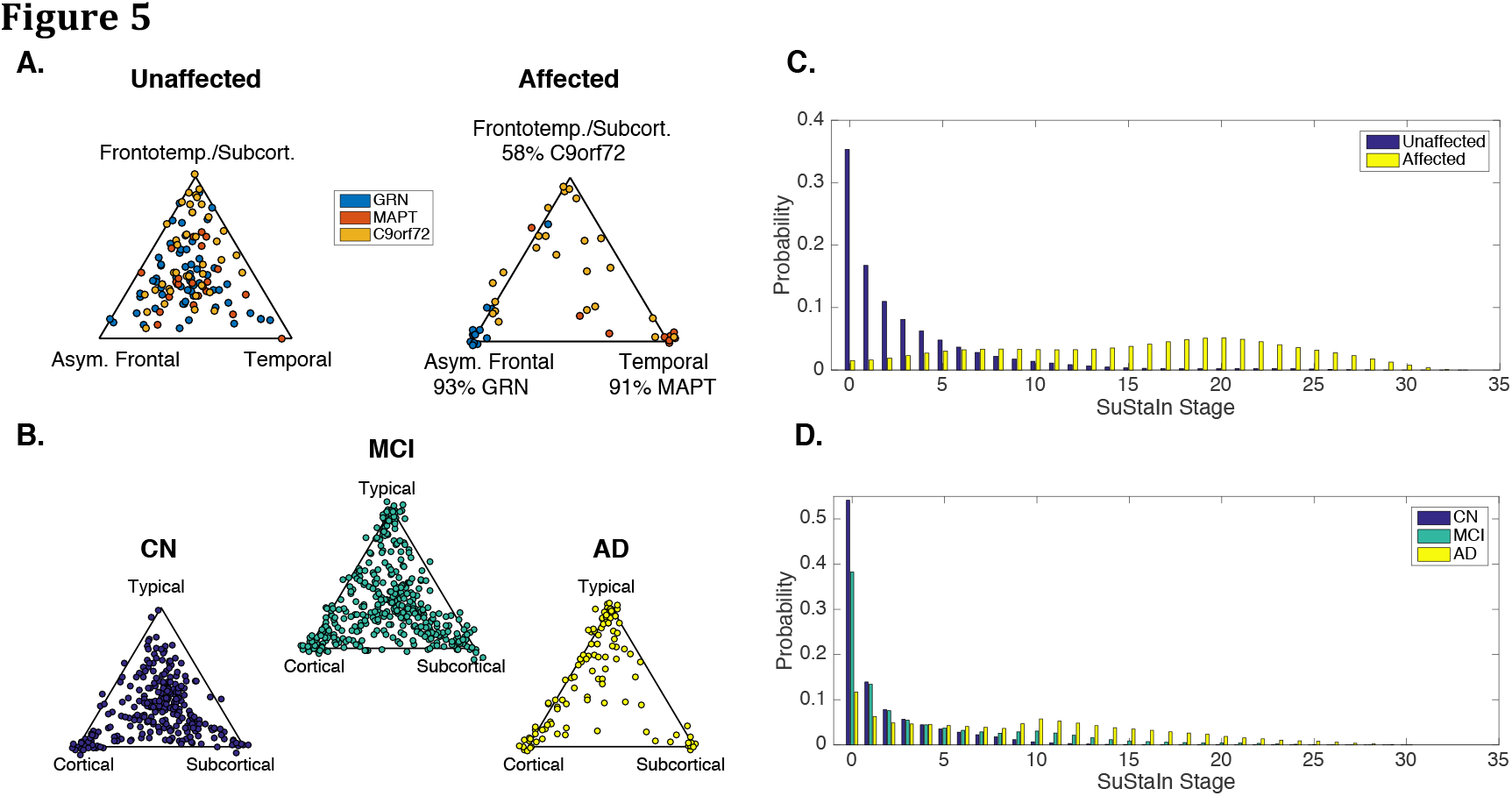
SuStaIn subtyping and staging of genetic frontotemporal dementia and Alzheimer’s disease. Subfigures A and B show the identifiability of the disease subtypes estimated by SuStaIn for genetic frontotemporal dementia, and Alzheimer’s disease. Each scatter plot visualises the probability that each individual belongs to each of the SuStaIn subtypes estimated for A. genetic frontotemporal dementia (as shown in Figure 2A), and B. Alzheimer’s disease (as shown in Figure 3). In the triangle scatter plots, each of the corners corresponds to a probability of 1 of belonging to that subtype, and 0 for the other subtypes; the centre point of the triangle corresponds to a probability of 1/3 of belonging to each subtype. Subfigures C and D show the probability subjects from each of the diagnostic groups belong to each of the SuStaIn stages for C. genetic frontotemporal dementia and D. Alzheimer’s disease.

**Figure 6.**
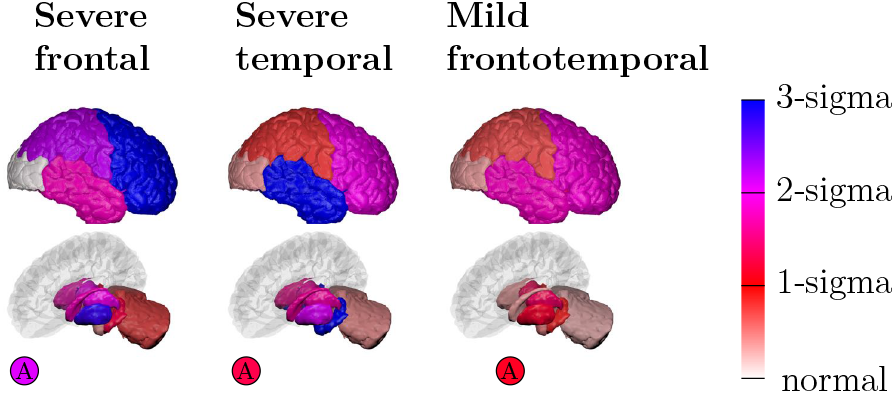
Subtypes-only model for GENFI; not accounting for disease stage heterogeneity. Brain diagrams as in Figure 2, but here each diagram represents a different subtype, which we refer to as severe frontal, severe temporal, and mild frontotemporal. There is no notion of disease stage in the subtypes-only model.

**Table 1.** A. SuStaIn (overall balanced accuracy 81%) Ability of subtypes to distinguish between different genetic types in symptomatic mutation carriers in GENFI using (A) SuStaIn subtypes (see Figure 2A) and (B) subtypes obtained from a subtypes-only model that accounts for heterogeneity in disease subtype but not disease stage (see Figure 6). Subfigures (C) and (D) show equivalent results when optimising SuStaIn and the subtypes-only model to distinguish genotype (see Methods: Classification of mutation groups using subtypes). Each entry is the percentage (number) of participants of a particular genetic type assigned to that subtype. The highlighted entries indicate the percentage (number) of participants assigned to the subtype that corresponds to the correct genetic type: *GRN* in blue, *MAPT* in red, and *C9orf72* in yellow. The results show that SuStaIn provides much better discrimination of the different genetic types than the subtypes-only model, demonstrating the added utility of a model that accounts for heterogeneity in disease stage.

|  | Asymmetric frontal | Temporal | Fronto-temporal | Subcortical |
| --- | --- | --- | --- | --- |
| GRN | 93% (13) | 0% (0) | 0% (0) | 7% (1) |
| MAPT | 9% (1) | 91% (10) | 0% (0) | 0% (0) |
| C9orf72 | 17% (4) | 25% (6) | 38% (9) | 21% (5) |

**B. Subtypes-only model (overall balanced accuracy 65%)**
|  | Severe Frontal | Severe Temporal | Mild Fronto-temporal |
| --- | --- | --- | --- |
| GRN | 64% (9) | 0% (0) | 36% (5) |
| MAPT | 9% (1) | 64% (7) | 27% (3) |
| C9orf72 | 17% (4) | 17% (4) | 67% (16) |

**C. Optimised SuStaIn (overall balanced accuracy 86%)**
| | Asymmetric frontal<br>(threshold $p > 0.65$) | Temporal<br>(threshold $p > 0.35$) | Fronto-temporal | Subcortical |
| --- | --- | --- | --- | --- |
| GRN | 93% (13) | 0% (0) | 0% (0) | 7% (1) |
| MAPT | 9% (1) | 91% (10) | 0% (0) | 0% (0) |
| C9orf72 | 4% (1) | 21% (5) | 42% (10) | 33% (8) |

**D. Optimised subtypes-only model (overall balanced accuracy 69%)**
| | Severe Frontal<br>(threshold $p > 0.99$) | Severe Temporal<br>(threshold $p > 0.99$) | Mild Fronto-temporal |
| --- | --- | --- | --- |
| GRN | 57% (8) | 0% (0) | 43% (6) |
| MAPT | 9% (1) | 64% (7) | 27% (3) |
| C9orf72 | 4% (1) | 8% (2) | 88% (21) |

**Table 2.** Utility of SuStaIn subtype and stage for predicting risk of conversion from mi d cognitive impairment to Alzheimer’s disease. Each row shows Hazards ratios for a different Cox Proportional Hazards model for predicting the risk of conversion from mild cognitive impairment to Alzheimer’s disease using ADNI data. Each column shows the estimated hazard ratio for each variable. Each hazards ratio tells you how the risk of conversion changes for each unit increase of a particular variable: a ratio of 1 means no modification of the risk, a ratio greater than 1 means there is an increase of the risk, and a ratio less than 1 means there is a reduction of the risk. For the first model (S-C-T) it is assumed that the hazard ratio increases multiplicatively from the Subcortical subtype (S) to the Cortical subtype (C) to the Typical subtype (T), i.e. the S-C-T model predicts that each SuStaIn subtype has a hazards ratio 1.57 times that of the previous subtype (i.e. the cortical group have a 1.57 times greater risk of conversion than the subcortical group, the typical group have a 1.57 times greater risk of conversion than the cortical group, and the typical group have a 2.46 (1.57^2^) times greater risk of conversion than the subcortical group). In the remaining models only two groups are compared at a time to demonstrate that the results are similar without this assumption, although the statistical power is reduced. Statistical significance is indicated as: ~ = p < 0.1, * = p < 0.05, ** = p < 0.01, † = p < 1 x 10^−3^. This result demonstrates the added utility of both disease subtypes and stages obtained from SuStaIn for predicting conversion between mild cognitive impairment and Alzheimer’s disease, with both subtype and stage modifying the risk of conversion.

|  | Model subtype | Model stage | Age | Sex | Education | APOE4 |
| --- | --- | --- | --- | --- | --- | --- |
| S-C-T | 1.57** | 1.13† | 0.98 | 0.98 | 0.93~ | 1.82† |
| S-C | 1.76~ | 1.16† | 0.95* | 1.03 | 0.92 | 1.53* |
| C-T | 1.48~ | 1.11† | 0.98 | 0.87 | 0.97 | 1.84† |
| S-T | 2.11* | 1.13† | 1.02 | 1.13 | 0.90* | 2.13† |

#### SuStaIn subtypes and stages are identifiable

Figure 5 illustrates the practical utility of SuStaIn’s disease subtyping and staging information for each neurodegenerative disease. Figure 5A shows that the identifiability of the SuStaIn subtypes in genetic FTD increases as the diseases progress, with the subtypes being strongly identifiable in symptomatic mutation carriers in GENFI. Figure 5B shows that the identifiability of the SuStaIn subtypes in AD also increases with disease progression, with a strong separation of the subtypes in ADNI participants with an AD diagnosis. The strong identifiability of the AD subtypes that SuStaIn achieves by accounting for temporal heterogeneity is in contrast to previous studies^23^ that model phenotypic but not temporal heterogeneity. Moreover, the identifiability is seen even at early disease stages (MCI), where many subjects cluster around the vertices of the triangles. Figures 5C and 5D show that the distribution of SuStaIn stages differs between diagnostic groups in both GENFI and ADNI, and provides a good separation of presymptomatic and symptomatic mutation carriers, and cognitively normal (CN) and AD.

#### SuStaIn subtypes discriminate FTD genotype

Table 1A shows the classification accuracy obtained using the SuStaIn subtypes in Figure 2 to discriminate the genotype of affected mutation carriers in GENFI. The SuStaIn subtypes give a balanced accuracy of 95% for the two-way classification task of distinguishing the homogeneous *GRN* and *MAPT* carrier groups. For the more challenging three-way classification task of distinguishing all genotypes in the presence of heterogeneity, the SuStaIn subtypes provide a balanced accuracy of 81%. A high proportion of the homogeneous *GRN* and *MAPT* carrier groups are correctly assigned to the asymmetric frontal lobe (93% of affected *GRN* carriers) and temporal lobe subtype progression patterns (91% of affected *MAPT* carriers). The heterogeneous *C9orf72* carrier group are much more difficult to classify, with a total of 58% of affected *C9orf72* carriers being assigned to the frontotemporal lobe and subcortical subtypes. Apart from heterogeneity, the *C9orf72* carriers are also more difficult to classify because the frontotemporal lobe and subcortical subtype progression patterns are more similar to the other subtypes; by evaluating the similarity of each pair of subtype progression patterns (see Methods: Similarity between two subtype progression patterns) we find that the asymmetric frontal lobe and temporal lobe subtypes have the most distinct progression patterns of any pair of subtypes; the asymmetric frontal lobe and frontotemporal lobe subtypes have the most similar progression patterns of any pair of subtypes.

The assignment to genotype in Table 1A is performed by simply allocating individuals to their most probable SuStaIn subtype and thus to the corresponding genotype. However, we can improve overall balanced accuracy with alternative choices of decision thresholds on the subtype probabilities that account for differences in the confidence with which individuals are assigned to groups (the homogeneous genetic groups are typically assigned to their corresponding phenotype with much higher confidence than the *C9orf72* group). Table 1C shows the classification accuracy obtained for discriminating genotype when the probability required for assignment to a particular subtype is optimised. After increasing the probability required for assignment to the asymmetric frontal lobe and temporal lobe subtypes corresponding to the homogeneous GRN and MAPT carrier groups, the overall balanced classification accuracy increases from 81% to 86%. The classification accuracy for assigning the homogeneous *GRN* and *MAPT* carrier groups to their corresponding phenotype remains high, with 93% of affected *GRN* carriers being assigned to the asymmetric frontal lobe subtype and 91% of affected *MAPT* carriers being assigned to the temporal lobe subtype. The assignment of *C9orf72* mutation carriers to their genotype improves substantially, with the proportion of affected *C9orf72* mutation carriers being assigned to the frontotemporal lobe and subcortical subtypes increasing from 58% to 75%. Of note is that the optimised threshold for assigning affected *MAPT* carriers to the temporal lobe subtype (probability of 0.35) is much lower than that for assigning affected *GRN* carriers to the asymmetric frontal lobe subtype (probability of 0.65). This is likely due to the presence of outliers in the *MAPT* group (Figure S1).

#### SuStaIn out-performs subtypes-only models for discriminating FTD genotype

Table 1B shows the classification accuracy obtained using a subtypes-only model (see Figure 6), which does not account for temporal heterogeneity, to discriminate the genotype of affected mutation carriers in GENFI. The SuStaIn subtypes out-perform the subtypes-only model. The subtypes-only model gives a balanced accuracy of 92% compared to 95% using SuStaIn for the two-way classification task of distinguishing *GRN* and *MAPT* carrier groups; the subtypes-only model gives a balanced accuracy of 65% compared to 81% using SuStaIn for the three-way classification task of distinguishing all genotypes. We further performed the same optimisation of the probability required to assign individuals to different subtypes for classification of genotype for the subtypes-only model (shown in Table 1D) as we did for SuStaIn (Table 1C). Again the SuStaIn subtypes substantially out-perform the subtypes-only model; the optimised subtypes-only model gives a balanced accuracy of 69% compared to 86% using SuStaIn. In the subtypes-only model the majority of misclassifications arise from the earlier stage affected *GRN* and *MAPT* carriers being assigned to the mild frontotemporal subtype associated with *C9orf72* carriers.

#### SuStaIn subtypes and stages have predictive utility in AD

Table 2 shows that the SuStaIn subtypes and stages have predictive utility for the risk of conversion between diagnostic categories in ADNI. By fitting a Cox Proportional Hazards model, we found significant effects of baseline SuStaIn subtype (p=2.44×10^−3^) and stage (p=8.76×10^−11^) on an individual’s risk of conversion from mild cognitive impairment (MCI) to AD. Of the SuStaIn subtypes, the subcortical subtype is associated with the lowest risk of conversion, whilst the typical subtype is associated with the highest risk of conversion. Table S1 shows that SuStaIn out-performs subtypes-only and stages-only models at predicting the risk of conversion between diagnostic categories in ADNI. By performing likelihood ratio tests comparing SuStaIn to subtypes-only and stages-only we find that SuStaIn provides a significantly better fit than both subtypes-only (p=3.96×10^−5^) and stages-only (p=7.18×10^−4^) models. This shows that both the subtypes and stages estimated by SuStaIn provide additional information for estimating the risk of conversion from MCI to AD.

## Discussion

In this study we introduce SuStaIn – a new and powerful tool for data-driven disease phenotype discovery, providing new insights into disease aetiology, and new power for patient stratification in clinical trials and healthcare. Using the GENFI dataset we are able to show that SuStaIn can successfully recover known distinct progression patterns in genetic FTD that correspond to individuals with different genotypes. SuStaIn further reveals within-group heterogeneity for carriers of a mutation in the *C9orf72* gene and characterises the heterogeneity as distinct temporal progression patterns in two subtypes. This demonstrates the utility of SuStaIn for data-driven disease phenotype discovery, and provides new biological insight into the *C9orf72* mutation. Application of SuStaIn to the 3T ADNI dataset provides data-driven support for post-mortem neuropathological findings, finding three distinct AD subtypes. These three subtypes are corroborated using a largely independent data set (ADNI 1.5T). The disease subtype characterisation SuStaIn provides goes much further than post-mortem neuropathological studies^3,28^, or other machine-learning techniques^23^, by characterising the temporal trajectory of each subtype, enabling *in vivo* classification of subjects by disease stage as well as disease subtype. We demonstrate the utility of SuStaIn for *in vivo* patient subtyping and staging in both genetic FTD and AD. In genetic FTD, we show that the SuStaIn neuroimaging subtypes can distinguish affected carriers belonging to different genetic groups with high classification accuracy. In AD, we demonstrate that the SuStaIn subtypes are identifiable, even at early disease stages (MCI), and that the SuStaIn subtypes and stages have added utility for predicting conversion between clinical diagnoses, beyond models that do not account for phenotypic heterogeneity (in disease subtype) or temporal heterogeneity (in disease stage).

### Subtype progression patterns

#### New insights into FTD heterogeneity

The asymmetric frontal lobe subtype and temporal lobe subtype in Figure 2 show clear similarities with previous studies of regional volume loss in *GRN* and *MAPT* mutation carriers respectively, i.e. asymmetric frontotemporoparietal lobe volume loss in *GRN* carriers and temporal lobe volume loss in *MAPT* carriers^24^. However, SuStaIn provides much greater detail and accuracy by avoiding reliance on crude *a priori* staging, e.g. via mean familial age of onset. The frontotemporal lobe subtype and subcortical subtype in Figure 2 both have features previously associated with *C9orf72* mutation carriers, e.g. widespread symmetric grey matter atrophy and volume loss in the cerebellum^24^, but SuStaIn assigns these features to two distinct disease subtypes, and further reveals the temporal progression of each subtype.

Several biological factors may produce the two subtypes observed in *C9orf72* mutation carriers, either individually or in combination. Clinically, whilst there is significant overlap, patients typically present with either a behavioural variant frontotemporal dementia or amyotrophic lateral sclerosis as their main phenotype^32^, and they can progress at various rates; genetically, the expansion length is variable and there are additional genetic modifiers (e.g. *TMEM106B* and *ATXN2*) that alter phenotype^33-35^; and pathologically, most cases have either type A or type B TDP-43 pathology^32^. Whilst further study is required to determine the biological factors that influence neuroanatomical phenotype, these findings demonstrate the power of SuStaIn in identifying hitherto unrecognised disease subtypes using clinical data, to generate hypotheses that can be tested using basic science approaches.

#### Outliers in the *MAPT* mutation carrier group

We also find evidence for the presence of outliers in the *MAPT* mutation carrier group, but numbers are too small to determine whether these outliers have a distinct progression pattern. We ran a post-hoc analysis to check for differences between the outliers and the general *MAPT* carrier population. We assigned *MAPT* mutation carriers to their most probable subtype of the two SuStaIn subtypes from fitting to the *MAPT* mutation carrier group alone. Among individuals with significant evidence of MRI atrophy (SuStaIn stage of greater than or equal to 5), four individuals were identified as outliers and nine as inliers. Although MAPT mutations have been commonly thought to have a very specific pattern of atrophy affecting the anterior and medial temporal lobes predominantly, one previous paper has shown that there can be a second pattern of atrophy in specific mutations, where the lateral temporal lobes are affected more than the medial regions^36^. Interestingly, the two pairs of individuals who constitute the outliers in our analysis all have P301L mutations, a mutation that falls into this second alternate atrophy pattern group in ^36^. None of the inliers in our analysis have P301L mutations, or V337M mutations, the other mutation identified in ^36^ as having an alternate atrophy pattern. This suggests that SuStaIn may be able to identify particular MAPT mutations that fall into this alternate group, but larger studies will be required to confirm this.

#### SuStaIn reveals the temporal progression of subtypes

In contrast to previous work, SuStaIn reveals the temporal progression of neurodegenerative subtypes, and is able to determine the optimal grouping into, and number of, subtypes supported by the data. In genetic FTD, we identify that the three genotypes are best described as four major phenotypes with distinct temporal progression patterns, with the *GRN* and *MAPT* mutation carrier groups each constituting a single major phenotype, but the *C9orf72* mutation carrier group best being described as two phenotypes. In AD, we find that there are three subtypes: a typical subtype for which atrophy starts in the hippocampus and amygdala; a cortical subtype for which atrophy begins in the nucleus accumbens, insula and cingulate; and a subcortical subtype for which atrophy originates in the pallidum, putamen, nucleus accumbens and caudate, with each subtype having its own distinct progression pattern. These temporal spreading patterns for distinct subtypes offer new biological insight. For example, the progression pattern of each subtype provides a view of how neurodegeneration spreads from a distinct origin over the rest of the brain that is uncorrupted by phenotypic heterogeneity. A key advantage of SuStaIn is that it provides a purely data-driven, hypothesis-free, reconstruction of the progression of neurodegenerative disease subtypes. However, these observations also have great potential to inform hypothesis-based mechanistic models^30,31^ of neurodegenerative disease, which explain the temporal progression of neurodegenerative diseases via various mechanisms of disease propagation over brain networks. This allows different hypothesised disease mechanisms to be evaluated on real data. Current mechanistic models implicitly assume a single disease progression pattern – an assumption often violated in disease data sets, but much more reasonable if focussed on particular SuStaIn subtypes.

#### Comparison of AD subtype progression patterns with neuropathological studies

Post-mortem histology^3^ and retrospectively-analysed MRI scans close to the time of death^28^ observe three distinct patterns of atrophy in late-stage AD patients: one focussed on the temporal lobe that is similar to the late stages of the typical SuStaIn subtype; one affecting predominantly cortical regions cf. late stages of the cortical SuStaIn subtype; and one with stronger subcortical involvement cf. late stages of the subcortical SuStaIn subtype. This gives confidence in the SuStaIn subtypes, which provide much greater information by revealing the progression of each subtype over time, including the earliest sites of regional volume loss. Moreover, and importantly for practical utility, the SuStaIn subtypes are identifiable *in vivo* using MRI.

#### Reproducibility of AD subtypes in an independent dataset

The three AD subtypes found in the 3T MRI data set are corroborated by the largely independent 1.5T MRI data set. However, there are some small differences between the subtype progression patterns of the three subtypes recovered in each data set. These differences are predominantly found in how early the nucleus accumbens begins to atrophy in the different subtypes: across all three subtypes the nucleus accumbens is found to atrophy earlier in the 3T subtypes than the 1.5T subtypes. A possible explanation for this is that the volume of the nucleus accumbens can be estimated more accurately using the higher field strength 3T MRI scans than the 1.5T MRI scans, and thus atrophy in the nucleus accumbens can be identified from an earlier stage in the 3T data set compared to the 1.5T data set.

#### AD outliers with a parietal subtype

In the 1.5T MRI data set we additionally find a small proportion (4%) of outliers with a parietal subtype. This small subgroup may represent outliers with a posterior cortical atrophy phenotype: comparing the Alzheimer’s disease Assessment Scale-cognitive subscale (ADAS-cog) scores between individuals with an AD diagnosis that are assigned to the parietal subgroup (N=6) and the typical AD subgroup (N=65), we find that the parietal subgroup have worse performance (Mann-Whitney U test) on certain praxic (Q6. Ideational Praxis, p=6.1x10^−3^, z=2.7) and spatially-demanding (Q14. Number Cancellation, p=4.9x10^−3^, z=2.8) subtests, but similar performance in memory domains (Q8. Word Recognition, p=0.81, z=-0.2; Q1. Word Recall, p=0.48, z=0.70). Additionally, the parietal subgroup is on average 10.3 years younger (Mann-Whitney U test, p=2.8x10^−3^ z=-3.0) than the typical AD subgroup.

### Disease subtyping and staging

#### Identifiability of SuStaIn subtypes

The previous study of Zhang et al. ^23^ also looked at the identifiability of AD subtypes using a subtypes-only model that does not account for temporal heterogeneity in disease stage. In contrast to the study of Zhang et al. ^23^, we observe strong identifiability of the subtypes in AD patients by accounting for heterogeneity in disease stage. This identifiability clearly increases with disease progression, with the subtypes being most identifiable in AD patients. However, even at early stages (MCI), many subjects cluster around the vertices of the triangles showing strong potential for identifying cohorts representative of each subtype.

#### Utility of SuStaIn subtypes and stages

SuStaIn shows strong capabilities for patient stratification in both genetic FTD and AD. SuStaIn provides high classification accuracy for differentiating the different mutation types in genetic FTD, and the AD subtypes are clearly identifiable. SuStaIn out-performs a subtypes-only model, giving a balanced classification accuracy of 81% for distinguishing genotype compared to 65% for the subtypes-only model. This provides compelling evidence that there is substantial heterogeneity in disease stage within different phenotypes, and that modelling this disease stage heterogeneity is important for better patient stratification. This is further demonstrated in AD, in which SuStaIn’s subtypes and stages substantially out-perform subtypes-only and stages-only models for predicting conversion between diagnostic categories. These early results are highly promising, particularly given that the particular choice of biomarkers used here (coarse regional brain volumes) is not optimised for stratification. Inclusion of a wider range of biomarkers in future will further improve the patient stratification provided by SuStaIn. For example in AD, incorporation of amyloid and neurofibrillary tangle measures, e.g. from amyloid and tau positron emission tomography (PET) scans, will enable stratification of individuals at the very earliest disease stages.

## Conclusion

We introduce SuStaIn – a new tool to disentangle and characterise the temporal and phenotypic heterogeneity of neurodegenerative diseases. We use it to uncover novel within-genotype phenotypes in genetic FTD and to characterise the temporal heterogeneity of both genetic FTD and AD subtypes with previously unseen detail. We further demonstrate SuStaIn’s potential as a patient stratification tool by showing that SuStaIn provides high classification accuracy for discriminating genotype in genetic FTD, as well as added utility for predicting conversion between clinical diagnoses in AD. SuStaIn has the potential to make substantial clinical impact as a tool for precision medicine and is readily applicable to any progressive disease, including other neurodegenerative diseases, lung diseases and cancer.

## Materials and Methods

### Data description

#### GENFI dataset

We used cross-sectional volumetric MRI data from GENFI (http://www.genfi.org.uk/). Subjects were included from the second data freeze of GENFI, which in total consisted of 365 participants recruited across 13 centres in the United Kingdom, Canada, Italy, Netherlands, Sweden, and Portugal. 313 had a usable volumetric T1-weighted MRI scan for analysis (15 participants did not have a scan and the other participants were excluded as the scans were of unsuitable quality due to motion, other imaging artefacts, or pathology unlikely to be attributed to frontotemporal dementia). The 313 participants included 141 non-carriers, 123 presymptomatic carriers, and 49 symptomatic carriers. Of the 123 presymptomatic mutation carriers there were 62 *GRN*, 39 *C9orf72*, and 22 *MAPT* carriers. Of the 49 symptomatic carriers, there were 14 *GRN*, 24 *C9orf72*, and 11 *MAPT* carriers. The acquisition and post-processing procedures for GENFI have been previously described in ^29^. Briefly, cortical and subcortical volumes were generated using a multi-atlas segmentation propagation approach^37^, combining cortical regions of interest to calculate grey matter volumes of the entire cortex, separated into the frontal, temporal, parietal, occipital, cingulate, and insula cortices. In addition to regional volumetric measures, we also included a measure of asymmetry, which is calculated as the absolute value of the difference between the volumes of the right and left hemispheres, normalised by the total volume of both hemispheres. This asymmetry measure was log transformed to improve normality.

#### ADNI dataset

Data used in the preparation of this article were obtained from the Alzheimer’s Disease Neuroimaging Initiative (ADNI) database (http://adni.loni.usc.edu). The ADNI was launched in 2003 by the National Institute on Aging (NIA), the National Institute of Biomedical Imaging and Bioengineering (NIBIB), the Food and Drug Administration (FDA), private pharmaceutical companies and non-profit organizations, as a $60 million, 5-year public-private partnership. For up-to-date information, see http://www.adni-info.org. Written consent was obtained from all participants, and the study was approved by the Institutional Review Board at each participating institution.

We downloaded data from Laboratory Of Neuro Imaging (LONI; http://adni.loni.usc.edu) on 11 May 2016 and constructed two cross-sectional volumetric MRI datasets for SuStaIn model fitting: those with higher (3T) and lower (1.5T) field strength. The inclusion criteria for the 3T and 1.5T datasets were having cross-sectional FreeSurfer volumes available that passed overall quality control from either a 3T (processed using FreeSurfer Version 5.1) or a 1.5T (processed using FreeSurfer Version 4.3) MRI scan. The 3T dataset consisted of 793 subjects (183 cognitively normal, 86 significant memory concern, 243 early mild cognitive impairment, 164 late mild cognitive impairment, 117 Alzheimer’s disease), of which 73 were enrolled in ADNI-1, 99 were enrolled in ADNI-GO, and 621 were enrolled in ADNI-2. The 1.5T dataset consisted of 576 ADNI-1 subjects (180 cognitively normal, 274 late mild cognitive impairment, 122 Alzheimer’s disease). The 1.5T and 3T datasets are largely independent: only 59 subjects (14 cognitively normal, 33 late mild cognitive impairment, 12 Alzheimer’s disease) have both 1.5T and 3T scans. We downloaded processed cross-sectional FreeSurfer volumes for 1.5T and 3T scans, using FreeSurfer Versions 4.3 and 5.1, and quality control ratings. We retained only the volumes that passed overall quality control, and normalised them by regressing against total intracranial volume. We further downloaded demographic information for covariate correction: age, sex, education, and APOE genotype from the ADNIMERGE table. We downloaded follow-up information to test the association of the SuStaIn model subtypes and stages with longitudinal outcomes, consisting of diagnostic follow-up data and cognitive test scores from the mini-mental state examination (MMSE). We also downloaded baseline CSF measurements of Aβ1-42, which we used to identify a control population.

#### Z-scores

We expressed each regional volume measurement as a z-score relative to a control population: in GENFI we used data from all non-carriers, in ADNI we used amyloid-negative cognitively normal subjects, defined as those with a CSF Aβ1-42 measurement greater than 192 pg/mL^38^. This gave us a control population of 48 amyloid-negative cognitively normal subjects for the 3T dataset, and 56 amyloid-negative cognitively normal subjects for the 1.5T dataset. We used these control populations to determine whether the effects of age, sex, education, or number of APOE4 alleles (ADNI only) were significant, and if so to regress them out. We then normalised each dataset relative to its control population, so that the control population had a mean of 0 and standard deviation of 1. Because regional brain volumes decrease over time the z-scores become negative with disease progression, so for simplicity we took the negative value of the z-scores so that the z-scores would increase as the brain volumes became more abnormal.

### Mathematical modelling

#### SuStaIn modelling

We formulate the model underlying SuStaIn as groups of subjects with distinct patterns of biomarker evolution (see Mathematical Model). We refer to a group of subjects with a particular biomarker progression pattern as a subtype. The biomarker evolution of each subgroup is described as a series of events, where each event corresponds to a biomarker reaching a particular z-score compared to a control group. This linear z-score event-based model is based on the event-based model in ^7,8,39^, but reformulates the events so that they represent the continuous linear accumulation of a biomarker from one z-score to another, rather than an instantaneous switch from a normal to an abnormal level. The resulting mixture of linear z-score event-based models describes the biomarker evolution of each subgroup as a piecewise linear trajectory, with a constant noise level that is derived from a control population (see Mathematical model). The model assumes a fixed number of subtypes C, for which we estimate the proportion of subjects *f* that belong to each subtype, and the order *Sc* in which biomarkers reach each z-score for each subtype *c = 1…C*. We determine the optimal number of subtypes *C* for a particular dataset through 10-fold cross-validation (see Cross-validation).

#### Mathematical model

The linear z-score event-based model underlying SuStaIn is based on a continuous generalisation of the original event-based model^7,8^, which we describe first.

The event-based model in ^7,8^ describes disease progression as a series of events, where each event corresponds to a biomarker transitioning from a normal to an abnormal level. The occurrence of an event, *E_i_*, for biomarker *i* = 1… *I*, is informed by the measurements *x_i_* of biomarker *i* in subject *j, j* = 1…*J*. The whole dataset *X* = {*x_ij_* | *i* = 1… *I, j* = 1…*J*} is the set of measurements of each biomarker in each subject. The most likely ordering of the events is the sequence *S* that maximises the data likelihood

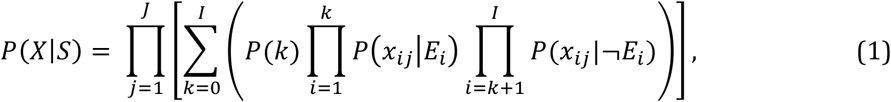

where *P*(*x*|*E_i_*) and *P*(*x*|¬*E_i_*) are the likelihoods of measurement *x* given that biomarker *i* has or has not become abnormal, respectively. *P*(*k*) is the prior likelihood of being at stage *k*, at which the events *E*_1_,…, *E_k_* have occurred, and the events *E*_*k*+1_,…, *E_I_* have yet to occur. The model uses a uniform prior, so that 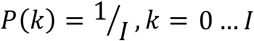. The likelihoods *P*(*x*|*E_i_*) and *P*(*x*|¬*E_j_* are modelled as normal distributions.

For the linear z-score event-based model we use in this work we reformulate the event-based model in (1) by replacing the instantaneous normal to abnormal events with events that represent the (much more biologically plausible) linear accumulation of a biomarker from one z-score to another. The linear z-score event-based model consists of a set of *N* z-score events *E_iz_*, which correspond to the linear increase of biomarker *i* = 1… *I* to a z-score *z_ir_* = *z_ir_*… *z_iR_i__*., i.e. each biomarker is associated with its own set of z-scores, and so *N* = Σ_*i*_ *R_i_*. Each biomarker also has an associated maximum z-score, *z_max_*, which it accumulates to at the end of stage *N*. We consider a continuous time axis, *t*, which we choose to go from *t* = 0 to *t* = 1 for simplicity (the scaling is arbitrary). At each disease stage *k*, which goes from 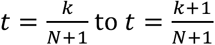, a z-score event *E_iz_* occurs. The biomarkers evolve as time *t* progresses according to a piecewise linear function *g_i_*(*t*), where

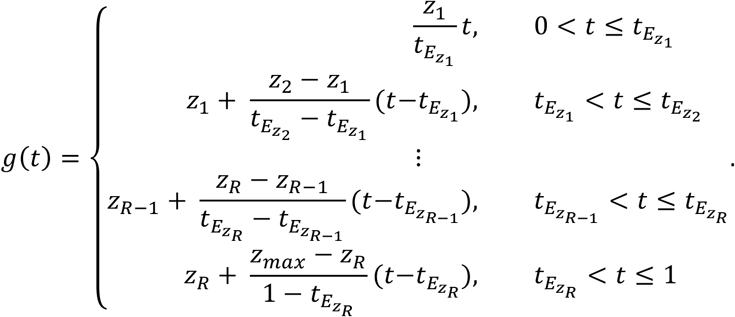

Thus, the times *t_E_iz__* are determined by the position of the z-score event *E_iz_* in the sequence *S*, so if event *E_iz_* occurs in position *k* in the sequence then 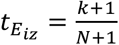.

To formulate the model likelihood for the linear z-score event-based model we replace (1) with

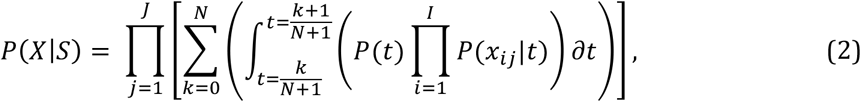

where,

NormPDF(*x, μ, σ*) is the normal probability distribution function, with mean *μ* and standard deviation *σ*, evaluated at *x*. We assume the prior on the disease time is uniform, i.e. *P*(*t*) = 1, as in the original event-based model.

The SuStaIn model is a mixture of linear z-score event-based models, hence we have

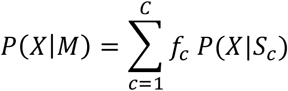

where *C* is the number of clusters (subtypes), *f* is the proportion of subjects assigned to a particular cluster (subtype), and *M* is the overall SuStaIn model.

#### Model fitting

We fit the SuStaIn model hierarchically by initialising the fitting of the *C* cluster (subtype) model from the *C-1* cluster model, i.e. we solve the clustering problem sequentially from *C = 1…C_max_*, where *C_max_* is the maximum number of clusters we would like to fit, initialising each model using the previous model. To fit the *C* cluster model using the *C-1* cluster model, we generate *C-1* candidate *C* cluster models by going through each of the *C-1* clusters in turn and finding their optimal split into two clusters; we then use this two cluster solution together with the other *C-2* clusters to initialise the fitting of the *C* cluster model. To optimise the *C* cluster model we use expectation maximisation, alternating between updating the sequences *S_c_* for each cluster and the fractions *f*_c_. Of these *C-1* candidate *C* cluster models, we choose the model with the highest likelihood as the solution to the clustering problem.

To find the optimal split of a cluster into two clusters, we initialise the assignments of data points to the two clusters randomly, find the optimal model parameters for these two data subsets, and use these cluster parameters to initialise the fitting of the two clusters. We repeat this procedure for 25 different start points (random cluster assignments) to find the maximum likelihood solution (see Convergence).

To find the optimal model parameters (the sequence *S* in which the biomarkers reach each z-score) for a single cluster we perform a greedy procedure whereby we initialise the sequence *S* randomly and then we go through each z-score event *e* in turn and find its optimal position in the sequence relative to the other z-score events, i.e. we fix the order of the subsequence *T = S\e* and evaluate the likelihood of the sequence in which the event *e* is placed at each possible position in the subsequence *T*. We keep updating the sequence *S* until convergence. Again we optimise the single cluster sequence *S* from 25 different random starting sequences to find the maximum likelihood solution (see Convergence).

#### Convergence

At several points in the model fitting we perform a greedy optimisation from a number of different starting points to find the maximum likelihood (the global optimum across the local optima reached from each start point) sequence or set of sequences. We find that the optimisation displays good convergence: all start points converge to a solution that is within a 1 × 10^−4^ % tolerance level (as a percentage of the maximum likelihood), and within the uncertainty estimated by the uncertainty estimation procedure (see Uncertainty estimation), meaning that each solution is sufficiently close to the maximum likelihood solution to be used for initialisation of the uncertainty estimation procedure.

#### Uncertainty estimation

In addition to estimating the most probable sequence *S_c_* for each subtype, we can determine the relative likelihood of all sequences for each subtype by evaluating the probability of each possible sequence. This gives us an estimate of the uncertainty in the ordering *S_c_*, which we summarise by plotting the probability that each z-score event appears at each position in the sequence for each subtype. We visualise this probability (see Figure 2 for example) using different colours to indicate the cumulative probability each region has reached a particular z-score: the cumulative probability of a region going from a z-score of 0-sigma to 1-sigma ranges from 0 in white to 1 in red, the cumulative probability of a region going from a z-score of 1-sigma to 2-sigma ranges from 0 in red to 1 in magenta, and the cumulative probability of a region going from a z-score of 2-sigma to 3-sigma ranges from 0 in magenta to 1 in blue. In practise the number of sequences is too large to evaluate all possible sequences so we use Markov Chain Monte Carlo (MCMC) sampling to provide an approximation to this uncertainty, as in ^7,8^. As in ^7,8^, we take 1,000,000 MCMC samples initialised from the maximum likelihood solution, checking that the MCMC trace shows good mixing properties.

#### Cross-validation

We performed 10-fold cross validation of the SuStaIn modelling results by dividing the data into 10 folds and re-fitting the model to each subset of the data, with one of the folds retained for testing each time. We report the consistency of the models across folds by computing the similarity between the progression patterns of two subtypes (see Similarity between two subtype progression patterns): the model fitted to each fold and the model fitted to the whole dataset. We evaluated the optimal number of subtypes using the Cross-Validation Information Criterion (CVIC)^40^, i.e. by evaluating the likelihood of each c-subtype model from *c = 1…C* on the test data for each fold and choosing the model with the highest out-of-sample likelihood *P*(*X*|*M*), or equivalently the lowest value of the CVIC, across all folds. The CVIC is defined as *CVIC* = –2 × log (*P*(*X*|*M*)), where *P*(*X*|*M*) is the probability of the data for a particular SuStaIn model, *M*, i.e. 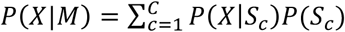. In cases where the evidence for a more complex model was not strong (a difference of less than 6 between the *CVIC* and the minimum *CVIC* across models, or equivalently a difference of less than 3 between the out-of-sample log-likelihood and the minimum out-of-sample log-likelihood across models), we favoured the less complex model to avoid over-fitting^41^.

#### Similarity between two subtype progression patterns

We measure the similarity of two subtype progression patterns using the Bhattacharyya coefficient^42^ between the position of each biomarker event in the two subtype progression patterns, averaged across biomarker events and MCMC samples. The Bhattacharyya coefficient measures the similarity of the distribution of the position of biomarker events in the subtype sequences and ranges from 0 (maximum dissimilarity) to 1 (maximum similarity).

#### Patient subtyping and staging

We assigned subjects to subtypes and stages predicted by the SuStaIn model by first evaluating the likelihood that they belonged to each subtype (by integrating over disease stage) and choosing the subtype with the highest likelihood, and then evaluating the probability they belonged to each stage of the most probable subtype and choosing the stage with the highest likelihood. When evaluating the likelihood we integrated over the set of MCMC samples to account for the uncertainty in the model parameters, rather than just evaluating the likelihood at the maximum likelihood parameters. This means that a patient’s model stage indicates the average position over the posterior distribution on the sequence given the data.

#### Comparison to subtypes-only and stages-only models

We compared our SuStaIn model to a subtypes-only model and a stages-only model. In the subtypes-only model, individuals are clustered together into groups based on the similarity of their biomarker measurements - without accounting for heterogeneity in disease stage. The stages-only model is a disease progression model where all subjects are assumed to be samples of a single common progression pattern - without accounting for heterogeneity in disease subtype. We formulated the subtypes-only and stages-only models so that they were as close as possible to the SuStaIn model, but did not model heterogeneity in disease stage or disease subtype, respectively. This allows us to assess the benefit of accounting for this disease stage or subtype heterogeneity in the SuStaIn model. The subtypes-only model consists of a mixture of Gaussians with unknown mean and variance. The subtypes-only model is fitted to symptomatic mutation carriers for GENFI, and Alzheimer’s disease subjects for ADNI, so that the subtypes correspond to a single diagnostic group. As done for the SuStaIn model, we evaluated the optimal number of clusters (subtypes) using the Cross-Validation Information Criterion^40^. The stages-only model is a special case of the SuStaIn model outlined in Mathematical Model, where only a single subtype is modelled.

#### Classification of mutation groups using subtypes

We performed two experiments to compare the ability of subtypes obtained from SuStaIn and the subtypes-only model to classify mutation carriers in GENFI into their different mutation groups. In the first experiment we simply assigned individuals to their most probable subtype and compared their assigned subtype with their mutation group. In the second experiment we optimised the probability required for assignment to each of the subtypes. This accounts for different amounts of heterogeneity within the different subtypes. In both experiments the classification results are reported as out-ofsample accuracies obtained through 10-fold cross-validation.

### Experiments

#### SuStaIn modelling of GENFI dataset

We applied SuStaIn modelling to various subgroups of the GENFI dataset: all 172 mutation carriers, 76 *GRN* mutation carriers, 63 *C9orf72* mutation carriers, 33 *MAPT* mutation carriers. For all the mutation carriers we fitted SuStaIn models of up to a maximum of 5 subtypes. For the *GRN, C9orf72* and *MAPT* mutation carriers we fitted SuStaIn models of up to a maximum of 3 subtypes. We chose the z-score events for the GENFI dataset to include z-scores of 1, 2, and 3 for each volume, but excluded z-score events where fewer than 10 mutation carriers had values that were greater than that z-score. The maximum z-score, which is reached at the final stage of the progression, was set to be 2, 3, or 5 depending on whether the maximum z-score event was 1, 2 or 3 respectively. We maintained the same z-score events across each of the GENFI experiments.

#### SuStaIn modelling of ADNI dataset

We applied SuStaIn modelling to the ADNI dataset, for which we tested SuStaIn models of up to a maximum of 5 subtypes. As for GENFI, we chose the z-score events to include z-scores of 1, 2, or 3 for each volume, but excluded z-score events where fewer than 10 subjects had values that were greater than that z-score. Again the maximum z-score, which is reached at the final stage of the progression, was set to be 2, 3, or 5 depending on whether the maximum z-score event was 1, 2 or 3 respectively.

## Acknowledgements

ALY is supported by a Doctoral Prize Fellowship from the EPSRC. NPO is supported by the Biomarkers Across Neurodegenerative Diseases program, which is funded by The Michael J. Fox Foundation for Parkinson’s Research, the Alzheimer’s Association, Alzheimer’s Research UK, and the Weston Brain Institute. DLT is supported by the UCL Leonard Wolfson Experimental Neurology Centre (PR/ylr/18575). KD is supported by an Alzheimer’s Society PhD Studentship. JMS acknowledges the support of the NIHR Queen Square Dementia BRU, the NIHR UCL/H Biomedical Research Centre, Wolfson Foundation, EPSRC (EP/J020990/1), MRC ( MR/L023784/1), ARUK (ARUK-Network 2012-6-ICE; ARUK-PG2017-1946; ARUK-PG2017-1946), Brain Research Trust (UCC14191) and European Union’s Horizon 2020 research and innovation programme (Grant 666992). JDR is supported by an MRC Clinician Scientist Fellowship (MR/M008525/1) and has received funding from the NIHR Rare Disease Translational Research Collaboration. The Dementia Research Centre is supported by Alzheimer’s Research UK, Brain Research Trust, and The Wolfson Foundation. This work was supported by the NIHR Queen Square Dementia Biomedical Research Unit and the NIHR UCL/H Biomedical Research Centre. This work is supported by EPSRC grants EP/J020990/01 and EP/M020533/1 and the *European Union’s Horizon 2020 research and innovation programme* under grant agreement No 666992 (EuroPOND: http://www.europond.eu).

